# Using visual encounter data to improve capture-recapture abundance estimates

**DOI:** 10.1101/2020.01.22.915314

**Authors:** Maxwell B. Joseph, Roland A. Knapp

## Abstract

Capture-recapture studies are widely used in ecology to estimate population sizes and demographic rates. In some capture-recapture studies, individuals may be visually encountered but not identified. For example, if individual identification is only possible upon capture and individuals escape capture, visual encounters can result in failed captures where individual identities are unknown. In such cases, the data consist of capture histories with known individual identities, and counts of failed captures for individuals with unknown identities. These failed captures are ignored in traditional capture-recapture analyses that require known individual identities. Here we show that if animals can be encountered at most once per sampling occasion, failed captures provide lower bounds on population size that can increase the precision of abundance estimates. Analytical results and simulations indicate that visual encounter data improve abundance estimates when capture probabilities are low, and when there are few repeat surveys. We present a hierarchical Bayesian approach for integrating failed captures and auxiliary encounter data in statistical capture-recapture models. This approach can be integrated with existing capture-recapture models, and may prove particularly useful for hard to capture species in data-limited settings.

## Introduction

Capture-recapture studies are widely used for estimating abundance and demographic rates, using information about the identities of captured individuals (Jolly 1965). This paper examines the case where individual identification requires capture, and identities of animals that are visually encountered but not captured are unknown. In such cases, “capture” is synonymous with “identification”. This often applies in capture-recapture studies of amphibians, where individual identification is only possible with the animal in hand. We also assume that if an individual escapes capture, it is not encountered again in the sampling occasion because it is hiding or otherwise inaccessible (Bailey and Nichols 2010, Joseph and Knapp 2018). Under these conditions, encounters leading to failed captures provide information about abundance, but this information is not readily used in traditional capture-recapture models.

When individuals can be encountered at most once on a survey, encounter and capture data both provide lower bounds on the total number of individuals in the population. Total abundance must be greater than or equal to the number of animals encountered in a survey. Similarly, total abundance must be greater than or equal to the number of unique individuals identified in the capture data. Capture data differ however, in that information accumulates over multiple surveys (Pollock 1982). For example, if two surveys occur on consecutive days in a closed population, then the total number of unique individuals captured across both surveys provides a lower bound on abundance.

Here, we show how visual encounter data can improve abundance estimates in capture-recapture studies for study designs where 1) individual identification requires capture, 2) a subset of encountered individuals are captured, and 3) individuals can be encountered at most once per sampling occasion. We develop a modified capture-recapture observation model, and investigate conditions under which encounter data improve abundance estimates. The methods presented here accommodate both failed captures and counts of animals collected separately from capture-recapture surveys, and can be integrated with existing capture-recapture models.

## Methods

### Model description

We adopt a hierarchical Bayesian approach in which an observation model depends on a state model, and both depend on some parameters (Berliner 1996). The state model describes the presence or absence of individuals in a population, and the observation model describes the visual encounter and capture process. The parameter model represents prior distributions for all remaining unknowns.

#### State model

Abundance can be estimated from capture-recapture data in a Bayesian framework using parameter-expanded data augmentation (Royle and Dorazio 2012). Here, *N*^∗^ unique individuals are observed, but *M > N*^∗^ individuals are modeled, augmenting the observed data with *M − N*^∗^ additional capture histories of animals that were never captured. The assumption is that the true abundance *N* is less than *M*, but greater than the number of observed individuals *N*^∗^.

Individuals *i* = 1, …, *M* are either “in the population” (*z*_*i*_ = 1) or not (*z*_*i*_ = 0), where the parameters *z*_1_, …, *z*_*M*_ are state parameters to be estimated, and abundance is *N* = Σ_*i*_ *z*_*i*_. These states can be modeled as conditionally independent Bernoulli random variables with probability parameter *ω*, where *ω* is the probability of an individual being in the population:

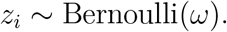

#### Observation model

On surveys *k* = 1, …, *K* an observer searches for individuals, encountering each individual with probability *η*. We assume individuals can only be encountered once at most. If an animal is encountered, it is captured with probability *κ*. Because individuals must be captured to be identified, encounter data are observed for captured individuals, but not for failed captures. Thus, encounter histories are partly observed. We assume the number of failed captures on each survey is observed.

Let 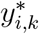 represent the categorical outcome for individual *i* on survey *k*. There are three possibilities (Figure 1):

1. The individual was not encountered 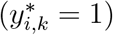, with probability *z*_*i*_(1 *− η*) + 1 *− z*_*i*_.
2. The individual was encountered but not captured 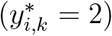, with probability *z*_*i*_*η*(1 *− κ*).
3. The individual was captured 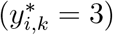, with probability *z*_*i*_*ηκ*.

**Figure 1.**
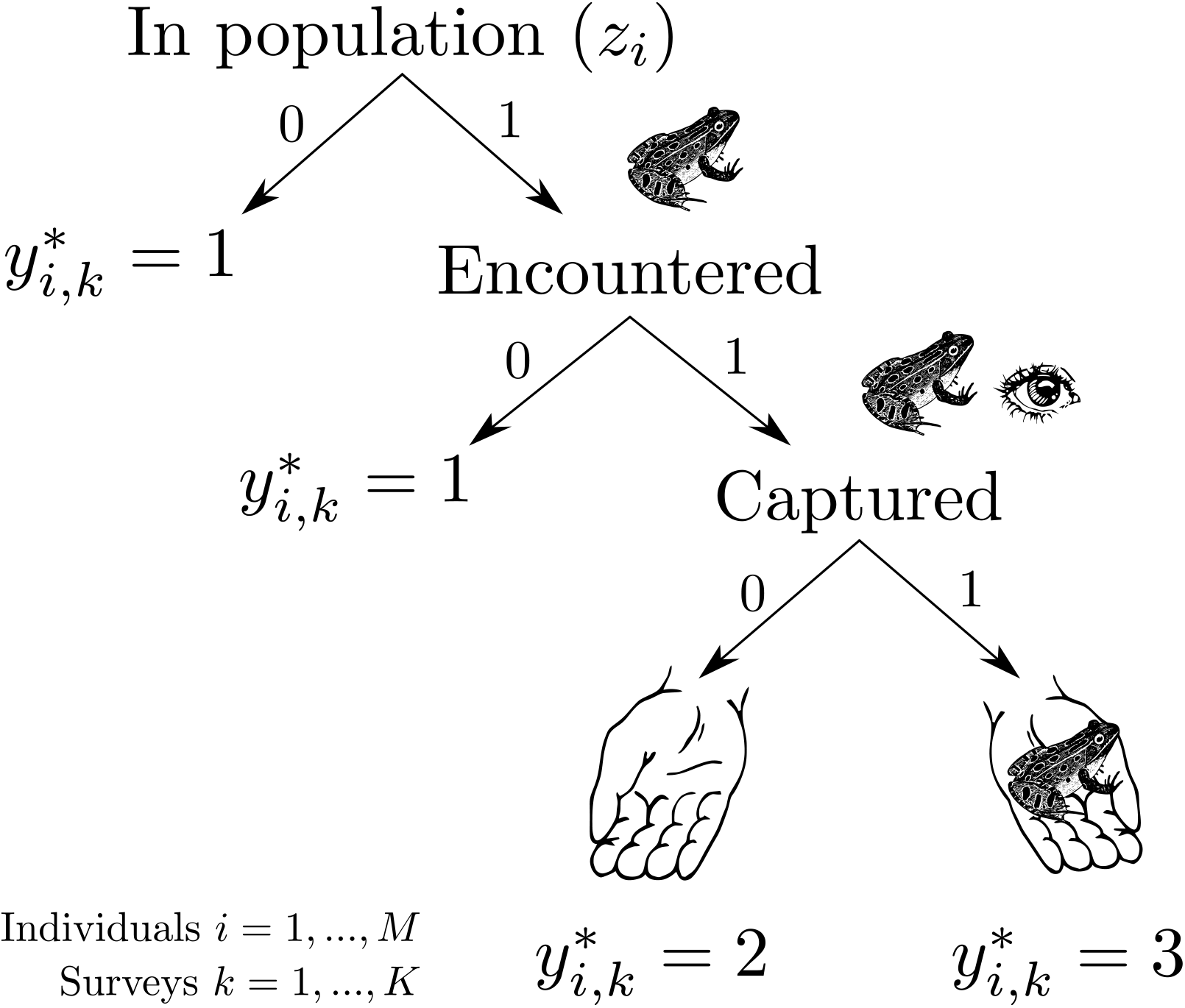
Conceptual diagram to represent the data model for individuals *i* = 1, …, *M* and sampling occasions or surveys *k* = 1, …, *K*. Each individual is either in the population (*z*_*i*_ = 1) or not (*z*_*i*_ = 0). Those that are not are never encountered. Those that are may be encountered on occasion *k* or not, and encountered individuals may or may not be captured and identified (we assume that identification requires capture, so that capture and identification are synonymous). Each path leads to a value of the partly observed quantity 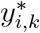, where 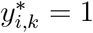 when animals are not encountered, 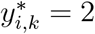 when animals are encountered but not identified, and 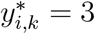 when animals are encountered and identified.

The first two outcomes are not observed. We observe a binary record of whether individual *i* was captured on survey *k*: *y*_*i,k*_, so that 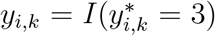, where *I* is an indicator function that is equal to one if the condition inside the parentheses is satisfied, otherwise it equals zero. In other words, 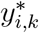 is observed only if 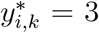. Additionally, the observed number of failed captures *f*_*k*_ corresponds to the sum 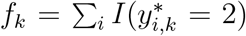. The observation model consists of two parts: one for the capture data:

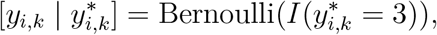

and another for the failed capture counts:

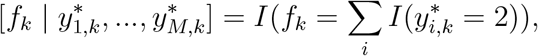

where square brackets denote a probability function. Alternatively, a “soft” constraint can be imposed as an approximation or to account for uncertainty in the number of failed captures:

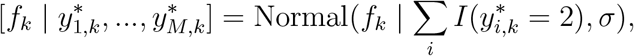

with *σ* set to some small fixed value.

#### Parameter model

The hierarchical model specification is completed by specifying prior distributions for remaining unknowns. Here, we use independent Uniform(0, 1) priors for all probabilities. This prior over the inclusion probability *ω* implies a discrete uniform prior for the true abundance from 0 to *M*.

#### Posterior distribution

The parameters consist of states *z*_1_, …, *z*_*M*_, outcomes 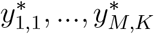, and the probabilities of inclusion (*ω*), encounter (*η*), and capture (*κ*). The data consist of the capture histories *y*_1,1_, …, *y*_*M,K*_, and counts of failed captures from each survey *f*_1_, …, *f*_*K*_. The posterior distribution of parameters given data, 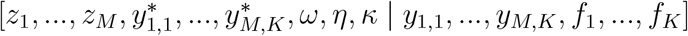, is proportional to:

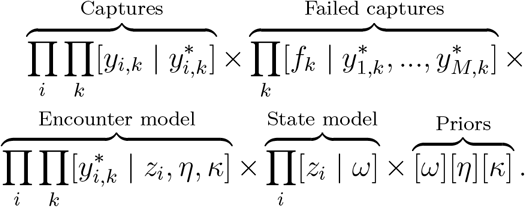

This model can be implemented in the popular BUGS language (Lunn et al. 2009), and we provide example specifications in Appendix S1.

### Auxiliary encounter data

In some cases, auxiliary encounter data are collected, such as during visual encounter surveys where individuals are counted on surveys, but no captures are attempted (Crump and Scott Jr 1994). Let *a*_*k*_ represent the number of unique individuals that were encountered on survey *k*. If encounters of individuals are conditionally independent, a binomial observation model provides a reasonable choice, where the number of trials is the population abundance *N* = Σ_*i*_*z*_*i*_ and the probability of success is the encounter probability *η*:

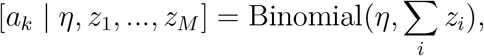

for surveys *k* = 1, …, *K*. This is the observation model used in N-mixture models (Royle 2004). In addition to potentially providing a higher lower bound on abundance, auxiliary encounters also provide additional information about encounter probabilities, because the encounter data are conditionally independent from encounters leading to captures. When combined with the capture-recapture model outlined above, the joint model of captures, failed captures, and auxiliary encounters comprise an integrated population model (Besbeas et al. 2002, Abadi et al. 2010). The posterior distribution is proportional to:

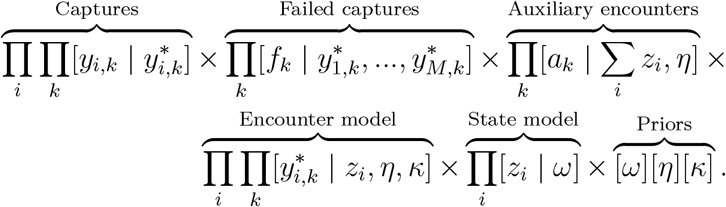

### Abundance lower bounds from encounter and capture data

The total population size is bounded from below by the number animals encountered on any one survey, assuming each animal can be encountered once at most (i.e., individuals are not double-counted). If *n*_*k*_ is the number of unique animals encountered on survey *k*, the lower bound on abundance from encounter data *n*_min_ = max(*n*_1_, …, *n*_*K*_) is the maximum of *K* independent binomial random variables with sample size *N* and probability *η*. The probability mass function of this lower bound is thus given by:

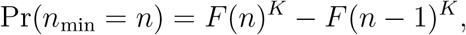

where *F* (*n*) is the cumulative distribution function of a binomial random variable. Population size must also be greater than or equal to the number of unique captured individuals. A probability mass function for the lower bound on abundance from capture data (*c*_min_: the number of unique captured individuals) can be derived with a binomial distribution. The binomial sample size is the true population size (*N*), and the probability of success is the probability of being captured one or more times: 1 *−* (1 *− ηκ*)^*K*^. The probability mass function for *c*_min_, the abundance lower bound derived from the capture data, is:

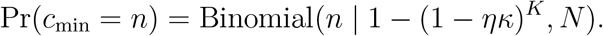

When the expected lower bound from encounter data exceeds the expected lower bound from capture data (𝔼(*n*_min_) *>* 𝔼(*c*_min_)), encounter data are expected to increase the precision of abundance estimates.

### Simulations

We empirically verified our theoretical results about the expected lower bounds on abundance provided by encounter and capture data using Monte Carlo simulation. We generated five replicate encounter-capture-recapture data sets for each parameter combination of *N* = 10, 50, 100, *K* = 3, 6, 9, and *η* and *κ* ranging from 0.01 to 0.99 in increments of 0.01, resulting in 441,045 unique data sets. For each parameter combination, we computed the empirical mean lower bounds from encounter and capture data, averaging over the five replicate iterations, and compared the results to the theoretical expectations generated from the probability mass functions for *n*_min_ and *c*_min_.

To understand the implications of bounding abundance for other parameters, we used a simulation study with known parameters across a range of repeat surveys (*K* = 3, *K* = 6, and *K* = 9). We visualized the joint posterior distribution of abundance and the probability of being encountered and captured, and compared these results to a simpler model: *M*_0_ - a capture-recapture model of a closed population with identical detection probabilities *p*_1_ = *…* = *p*_*K*_ = *p*, which ignores encounters that do not lead to captures (Royle and Dorazio 2008). This model can only estimate the marginal probability of capture *p* = *ηκ*. The observation model is *y*_*i,k*_ ∼ Bernoulli(*z*_*i*_*p*) for individuals *i* = 1, …, *M* on survey *k* = 1, …, *K*. The state model for *z* is unchanged, and uniform priors over (0, 1) were assigned to *ω* and *p*.

Draws from the posterior distributions of all models were simulated using JAGS, with six parallel Markov chain Monte Carlo chains, and 400,000 iterations per chain with an adaptation period of 200,000, a burn-in period of 40,000, and posterior thinning by 400 to reduce memory usage (Plummer and others 2003). Convergence was assessed using visual inspection of traceplots, and the potential scale reduction factor 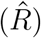 statistic (Gelman et al. 1992). All code to replicate the analyses is available in a research compendium at https://github.com/mbjoseph/secmr.

## Results

Across a range of abundances, encounter data are expected to provide a higher lower bound on abundance when capture probabilities are low, and when there are few repeat surveys (Figure 2). The boundary in the bivariate encounter-capture parameter space delineating the region where 𝔼(*c*_min_ *− n*_min_) *<* 0 shifts toward lower capture probabilities as the number of repeat surveys increases, and as abundance increases. Empirical average bounds from simulated capture-recapture data were in agreement with the theoretical expectations derived from the probability mass functions of *n*_min_ and *c*_min_ (Figure 3).

**Figure 2.**
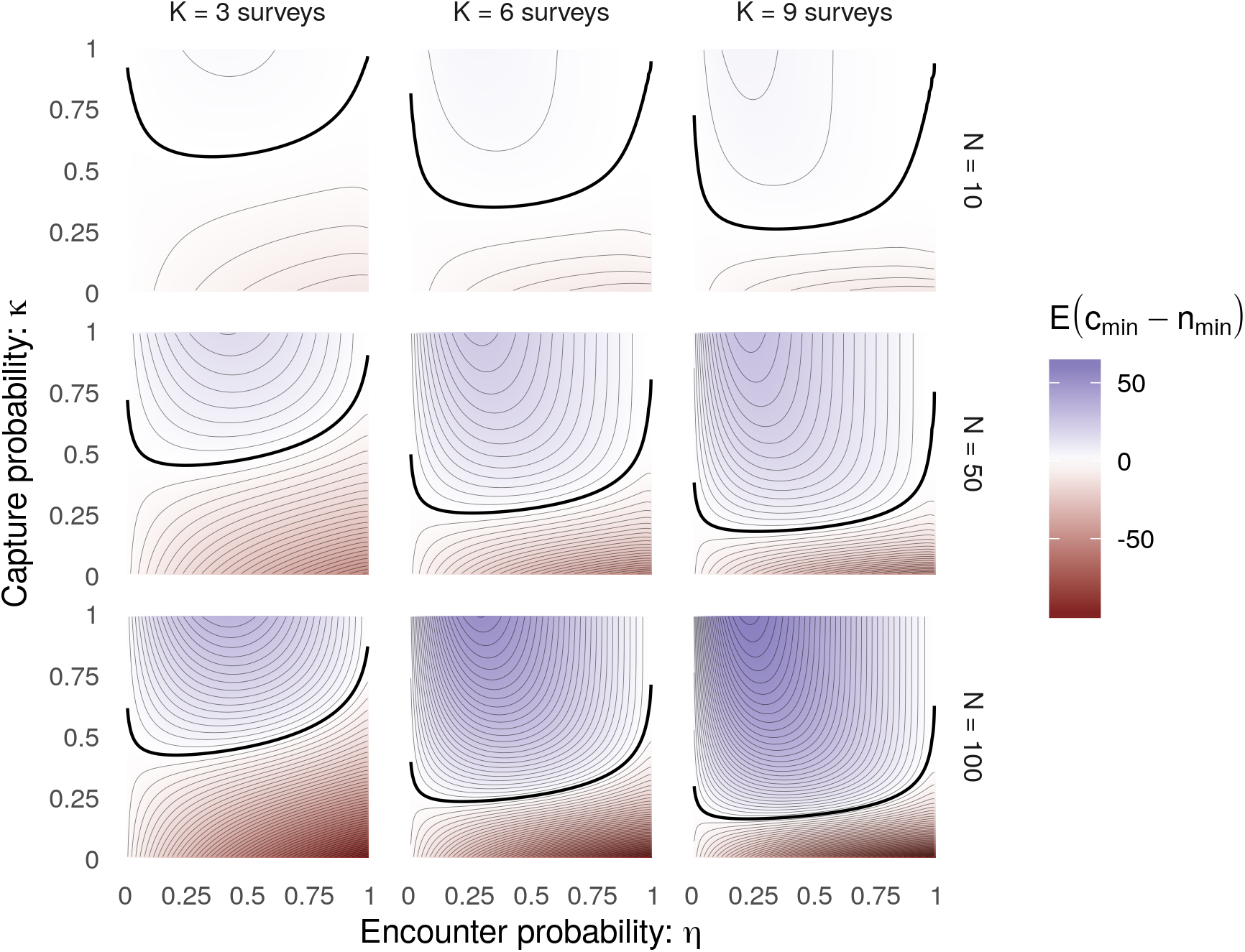
Expectations for the difference in abundance lower bounds provided by capture and encounter data as a function of the number of surveys *K*, abundance *N*, the encounter probability *η*, and the probability of capture conditional on an encounter *κ*. When the surface is red, encounter data are expected to increase the precision of abundance estimates by increasing the lower bound on true abundance. The heavy black line marks the null isocline where the expected difference is zero. Lighter lines represent contours spaced by 2 individuals.

**Figure 3.**
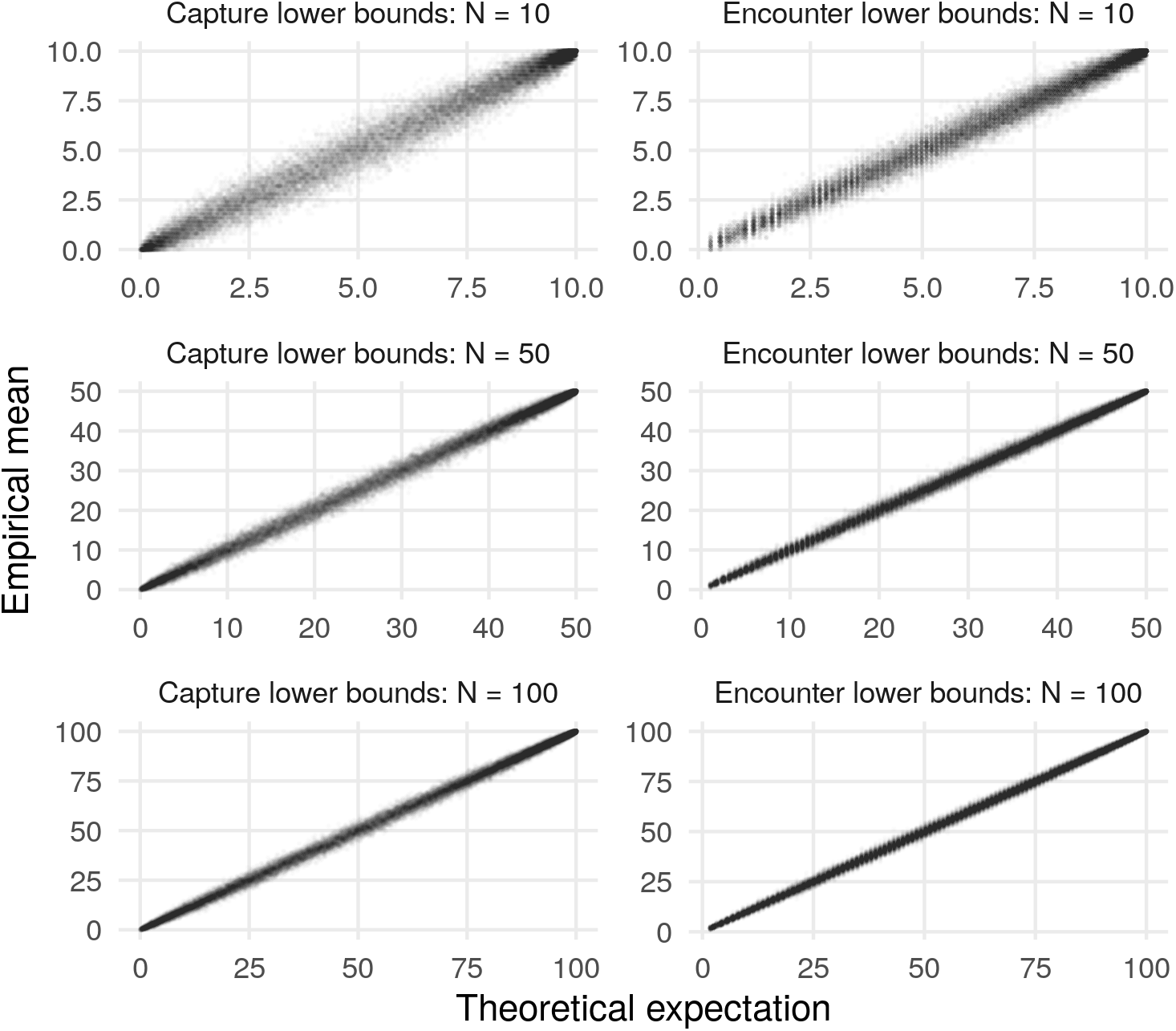
Empirical verification of theoretical expectations for lower bounds on abundance provided by encounter and capture data. Each point represents the empirical average of five replicate simulations across a range of parameter values, with panels separated by population size (*N*) and whether the lower bounds are derived from capture data (*c*_min_) or encounter data (*n*_min_).

When the lower bound on abundance is greater for encounter data than capture data, the joint model of encounters and captures produces a more precise estimate of abundance because there is zero probability mass below the lower bound on abundance. Furthermore, if there is posterior correlation between abundance and another parameter, encounter data can also increase the precision of the correlated parameter. For example, the marginal probability of capture and abundance are correlated in the posterior, so that increased posterior precision for abundance implies increased posterior precision of marginal capture probability (Figure 4).

**Figure 4.**
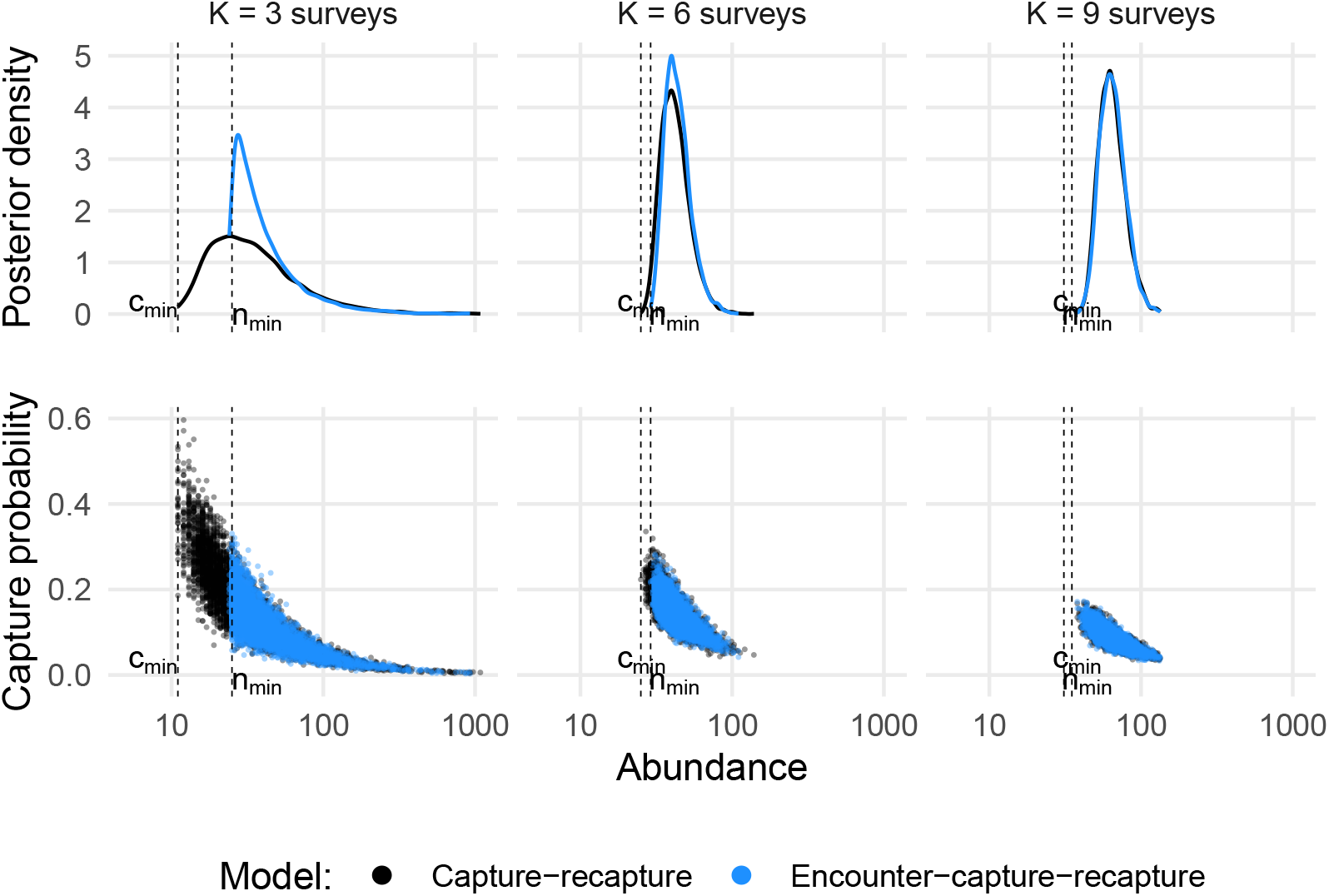
Samples from the posterior distribution of abundance (N, x-axis) and marginal capture probability *p* = *ηκ* for an individual in the population (y-axis). Each point is a sample from the posterior. Black points correspond to the baseline capture-recapture model *M*_0_, which does not include encounter data. Blue points correspond to an encounter-capture-recapture model that uses encounter data to bound abundance. Vertical dashed lines are shown for the lower bounds on abundance derived from encounter (*n*_min_) and capture (*c*_min_) data.

## Discussion

Failed captures and auxiliary encounters are likely to be most useful in capture-recapture studies when animals are hard to capture and the number of surveys is small. This expectation holds across a range of population abundance and encounter probabilities. In such cases, encounter data increase the precision of population abundance estimates by increasing the lower bound on abundance. Such data are essentially “free” in encounter-capture-recapture study designs, and can be included by modifying the likelihood function (and not the underlying state model) of capture-recapture models.

In addition to increasing the precision of abundance estimates, encounter data can increase the precision of parameter estimates for parameters that are correlated with abundance in the posterior distribution. For the simple model presented here, this includes the detection and inclusion probabilities. For more complex models that allow state evolution through time, this might include survival and recruitment probabilities. Thus, we expect that encounter data in general might provide information about abundance, and parameters relating to abundance and the measurement process.

These results are consistent with related findings for mark-resight studies where marked individuals are subject to incomplete identification. In such studies, accounting for failed identifications of marked individuals – analogous to failed captures – is most advantageous when identification probabilities are low – analogous to capture probabilities being low (McClintock et al. 2014b). However, in the encounter-capture-recapture scenario considered here, whether an animal is marked or not is unknown until it is captured, as would be the case for subdermal passive integrated transponder tags in an amphibian (Gibbons and Andrews 2004).

Here we assumed that individuals could be encountered at most once per sampling occasion. In other words, there is no double counting: each failed capture corresponds to one unique individual. The observation model developed here is not robust to violations of this assumption, because the total number of encounters sets a lower bound on the true population size. As a consequence, if individuals escape capture multiple times in the sampling sampling occasion, it is possible that the posterior for abundance might be misleadingly precise (i.e., the lower bound on population size would be too high). Therefore, we do not recommend this approach if individuals might be encountered multiple times on the same sampling occasion. This assumption is likely to hold for example in capture-recapture studies of amphibians in high elevation lakes, where individual animals that escape capture hide afterwards, e.g., underwater where they cannot be seen or captured (Joseph and Knapp 2018). If individuals are captured multiple times in the same sampling occasion, then it will be clear that the assumption of at most one encounter has been violated. In such cases, alternative encounter models may be necessary, e.g., a Poisson model that allows repeat encounters on a sampling occasion (Royle et al. 2009).

The model developed here relates to other approaches for handling imperfect individual identification in capture-recapture studies including misidentification (Link et al. 2010, McClintock et al. 2014a, Schofield and Bonner 2015) and partial identification (Augustine et al. 2018), and also to approaches that account for latent encounter histories with observed summary counts (Chandler et al. 2013). In particular, this approach might be viewed as an aspatial analog of the spatial model presented in Chandler et al. (2013) in which a subset of individual identities are available, with a Bernoulli (instead of a Poisson) encounter model, and a stochastic second stage “identification upon encounter” component. This approach can also be seen as a degenerate case of a partial identification model, in which there are no spatial (Augustine et al. 2018) or genetic data (Wright et al. 2009) available to inform the identities of individuals that have escaped capture.

Looking ahead, there are opportunities to build upon this approach. First, in terms of implementation, marginalization over the discrete latent variables might allow more efficient sampling from the posterior distribution. Second, because this model includes separate parameters for encounter and capture probabilities, covariates can be included separately for each of these components. This could be useful for example to account for predator avoidance behavior that might influence capture probabilities, and weather conditions that might influence encounter probabilities. Observer effects provide an additional use case: some observers might be better than others at finding or capturing individual animals.

In this paper, we presented a motivation for including encounter data in capture-recapture studies based on abundance lower bounds from encounter and capture data. Given that encounter data are included via a modified likelihood and not a modified state model, this approach can be readily integrated with a variety of capture-recapture models, and may be useful for hard to capture species in data-limited settings.

## Acknowledgements

This work was motivated by years experience in the field capturing (and failing to capture) amphibians in the Sierra Nevada and the Klamath mountains, and funded by a grant from the Yosemite Conservancy. We thank Ben Augustine for helpful comments on an earlier version of the manuscript.

## Appendix S1

This appendix includes model specifications and BUGS/JAGS code for failed capture and auxiliary encounter models. These models differ in the observation process, but the model of presence/absence states is identical. Each individual *i* = 1, …, *M* is either in the population (*z*_*i*_ = 1) or not (*z*_*i*_ = 0), where *z*_*i*_ ∼ Bernoulli(*ω*), and *ω* is an inclusion probability parameter. Abundance is the sum of these values *N* = Σ_*i*_*z*_*i*_.

### Baseline model

The baseline model is *M*_0_ and assumes equal detection probabilities of individuals *i* = 1, …, *M* on surveys *k* = 1, …, *K*, where *y*_*i,k*_ = 1 represents a capture of individual *i* on survey *k*, and *y*_*i,k*_ = 0 indicates no capture. If *p* is the capture probability, then *y*_*i,k*_ ∼ Bernoulli(*z*_*i*_*p*). The posterior distribution is:

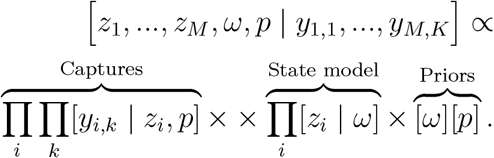

JAGS code for this model with uniform priors over *ω* and *p* is:

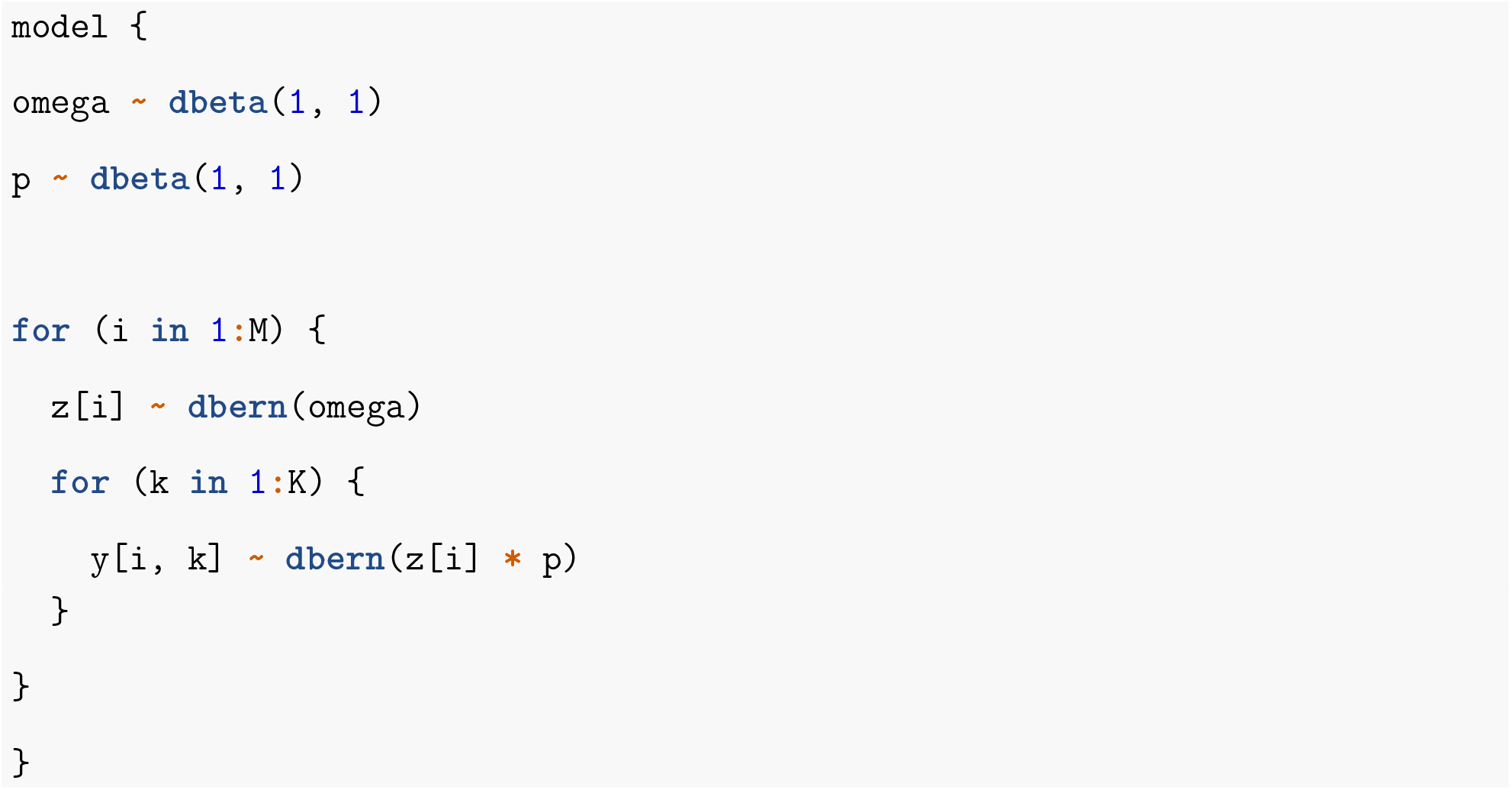

### Baseline model with failed captures

We add failed captures to the baseline model by introducing a categorical parameter 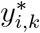, which represents “not encountered” 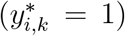, “failed capture” 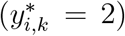, or “capture” 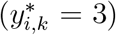. If *η* is the encounter probability, *κ* is the probability of capture conditional on encounter, and *f*_*k*_ is the number of failed captures, then as described in the main text the posterior is:

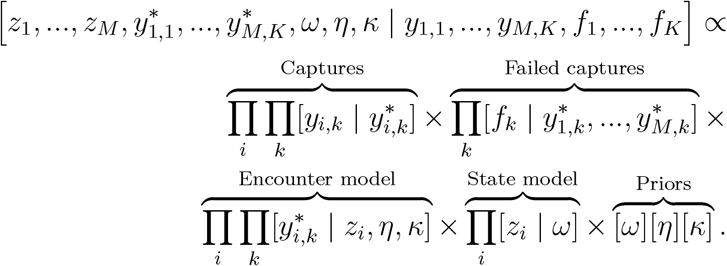

JAGS code for this model with uniform priors over *ω*, *η*, and *κ* is:

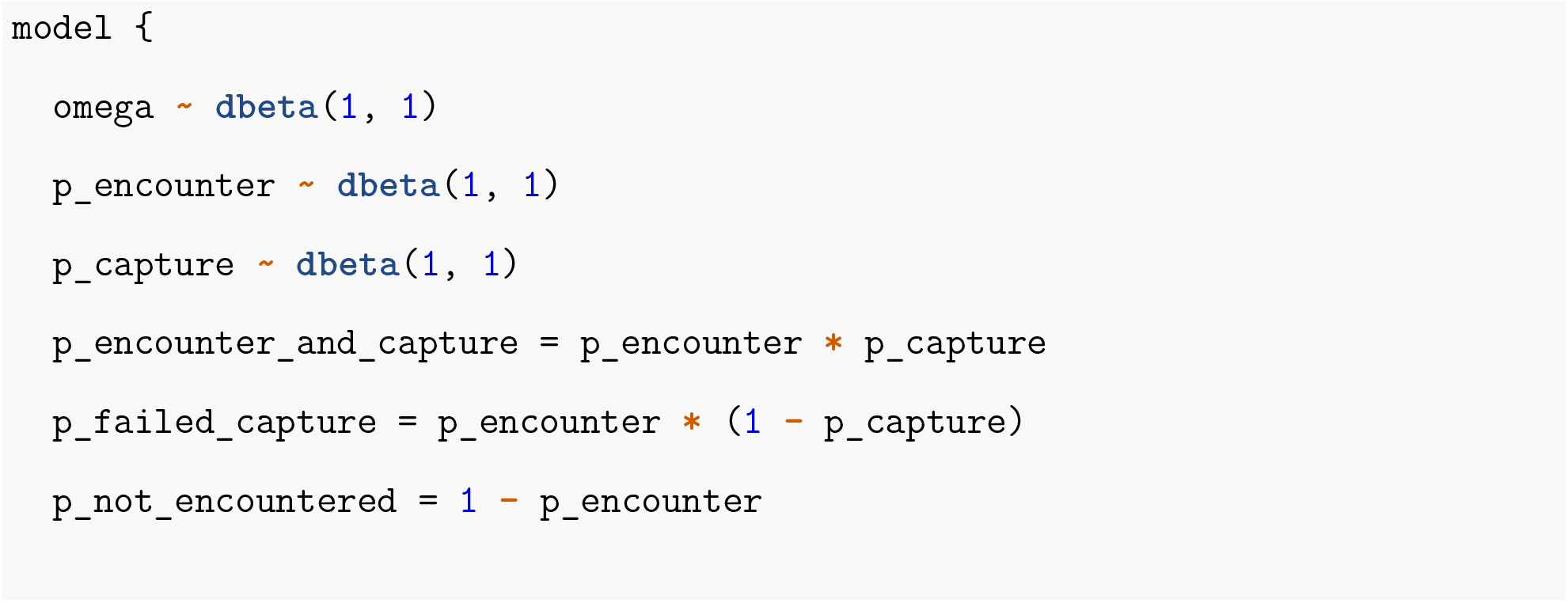
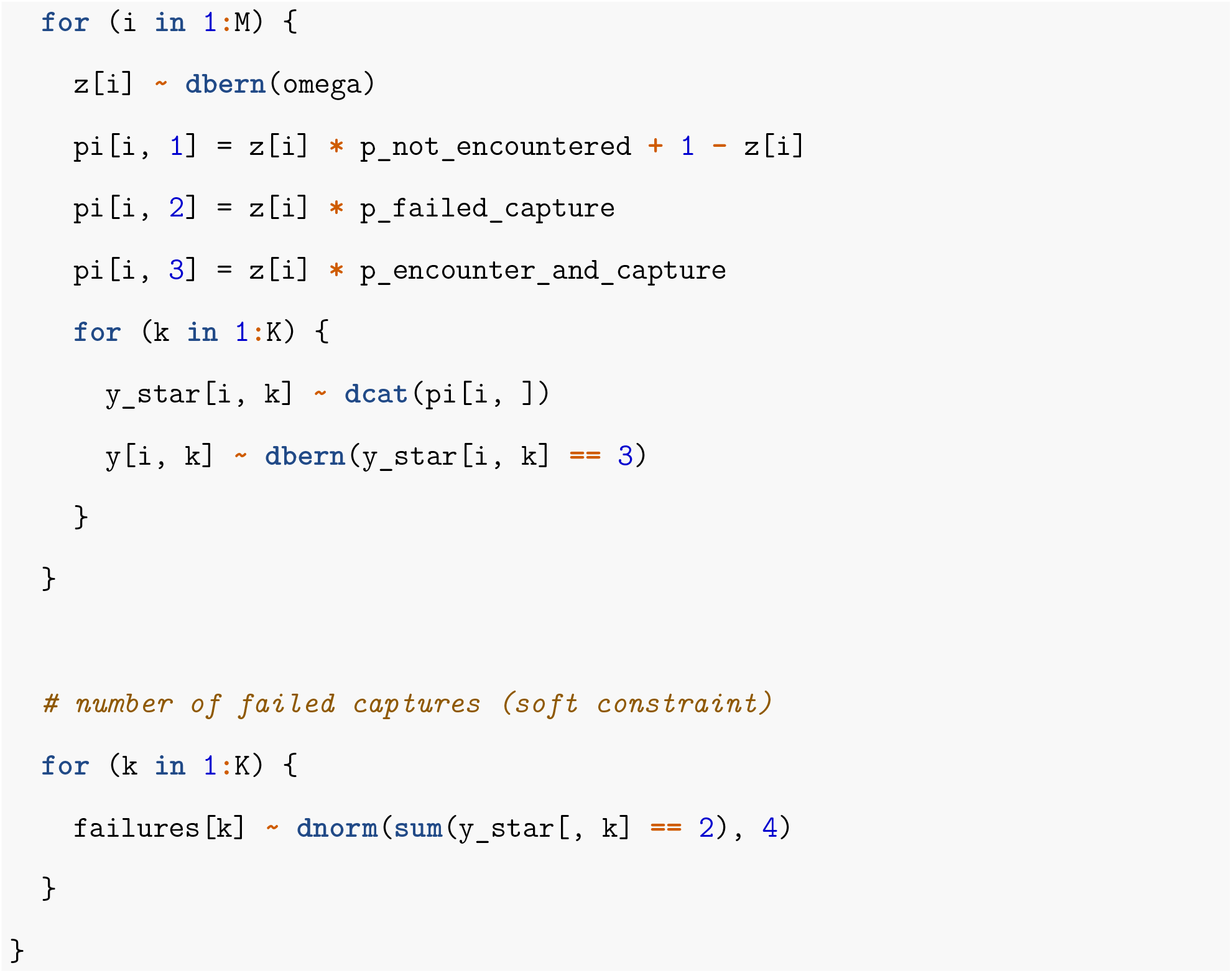

Note that the normal distribution for failed captures imposes a “soft” constraint on the sum 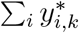. By adjusting the normal precision parameter, this constraint can be relaxed or made more strict. In the strictest case a hard constraint can be imposed via the “ones trick”, where a vector of ones is passed as data, and the likelihood for failed captures is represented as `ones[k] ~ dbern(sum(y_star[, k]) == failures[k])`.

### Baseline model with auxiliary encounters

To add auxiliary encounter data to the baseline model, we use the following likelihood for the number of auxiliary encounters *a*_*k*_ in survey *k* conditional on the true abundance *N* and the encounter probability *η*:

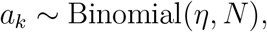

where *N* = Σ_*i*_*z*_*i*_. Note that there need not be an equal number (*K*) capture-recapture surveys and auxiliary encounter surveys, but here we assume this is the case to simplify notation. This model permits decomposing the parameter *p*, which represents the marginal probability of capture, into the product of the encounter probability *η* and the probability of capture conditional on encounter *κ*.

Then, the posterior is:

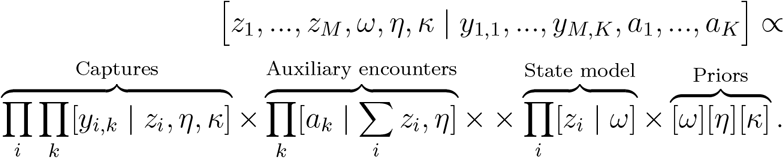

JAGS code for this model with uniform priors over *ω*, *η*, and *κ* is:

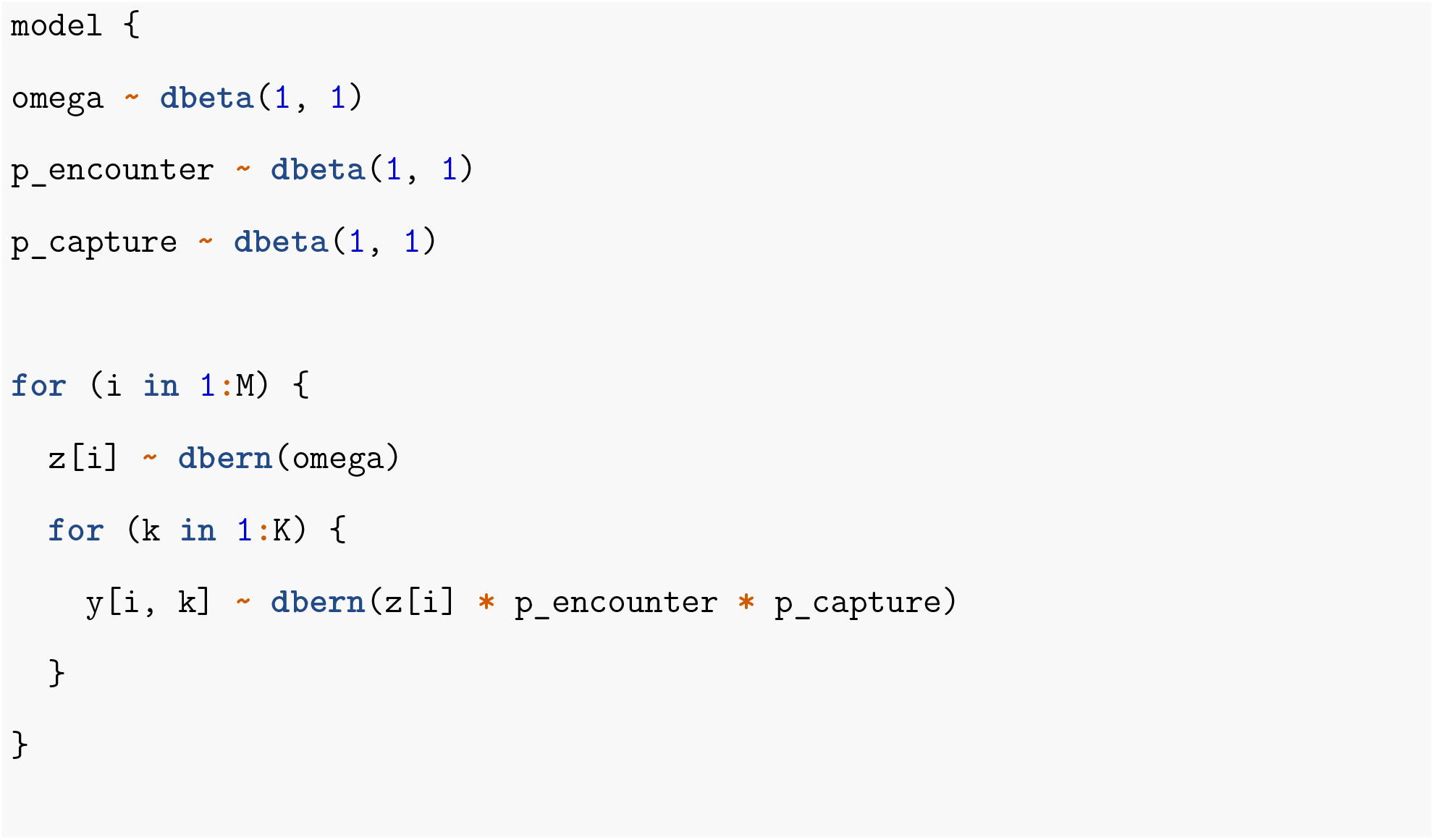
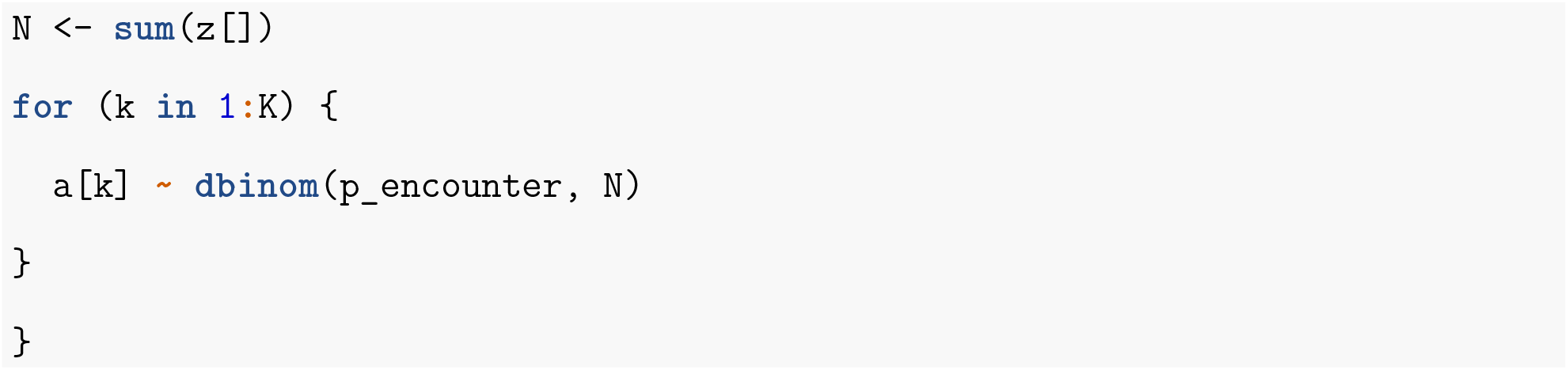

### Baseline model with failed captures and auxiliary encounters

Combining failed captures and auxiliary encounters includes the N-mixture model likelihood for auxiliary encounters in the failed capture model. As described in the main text, the posterior is:

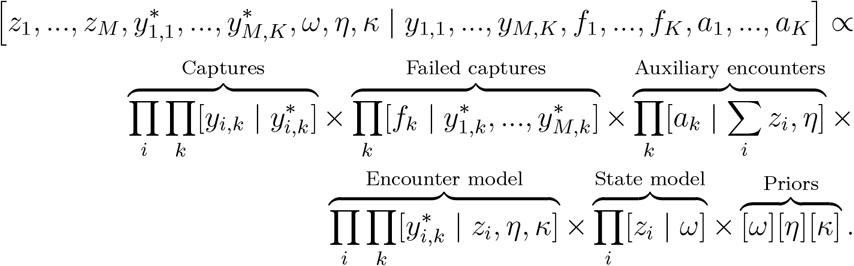

JAGS code for this model with uniform priors over *ω*, *η*, and *κ* is:

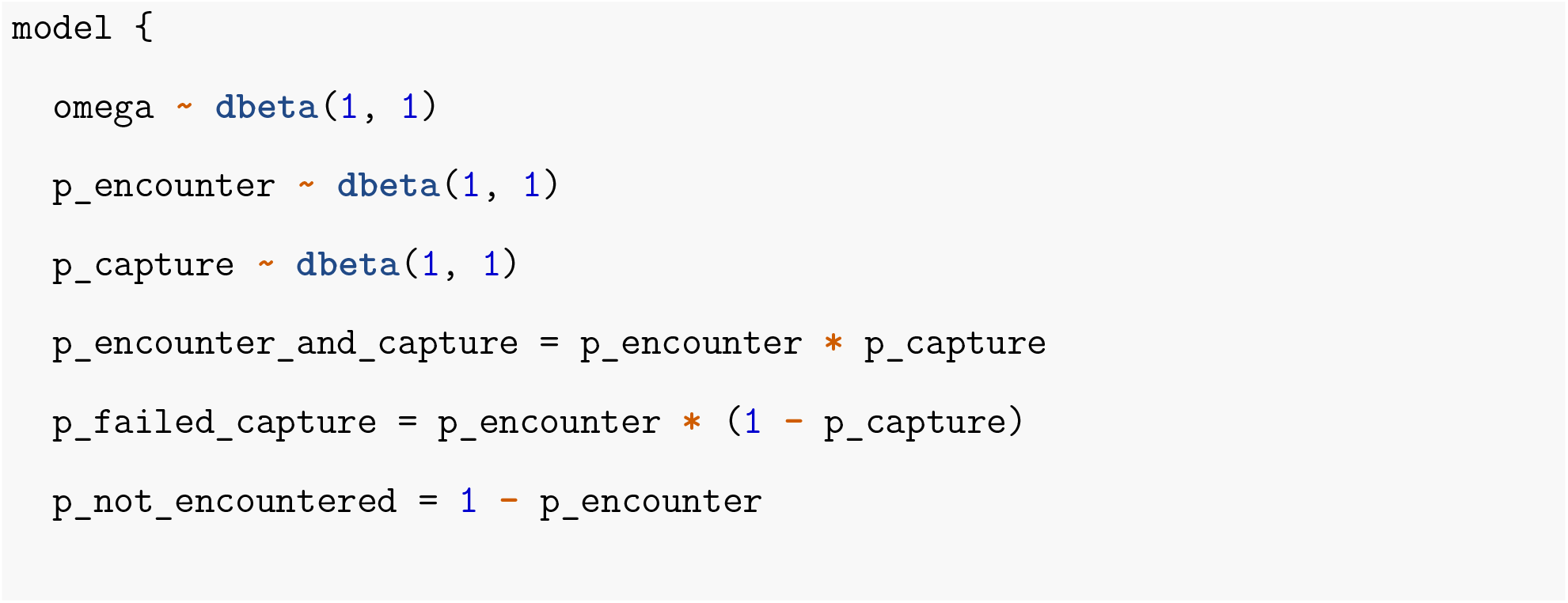
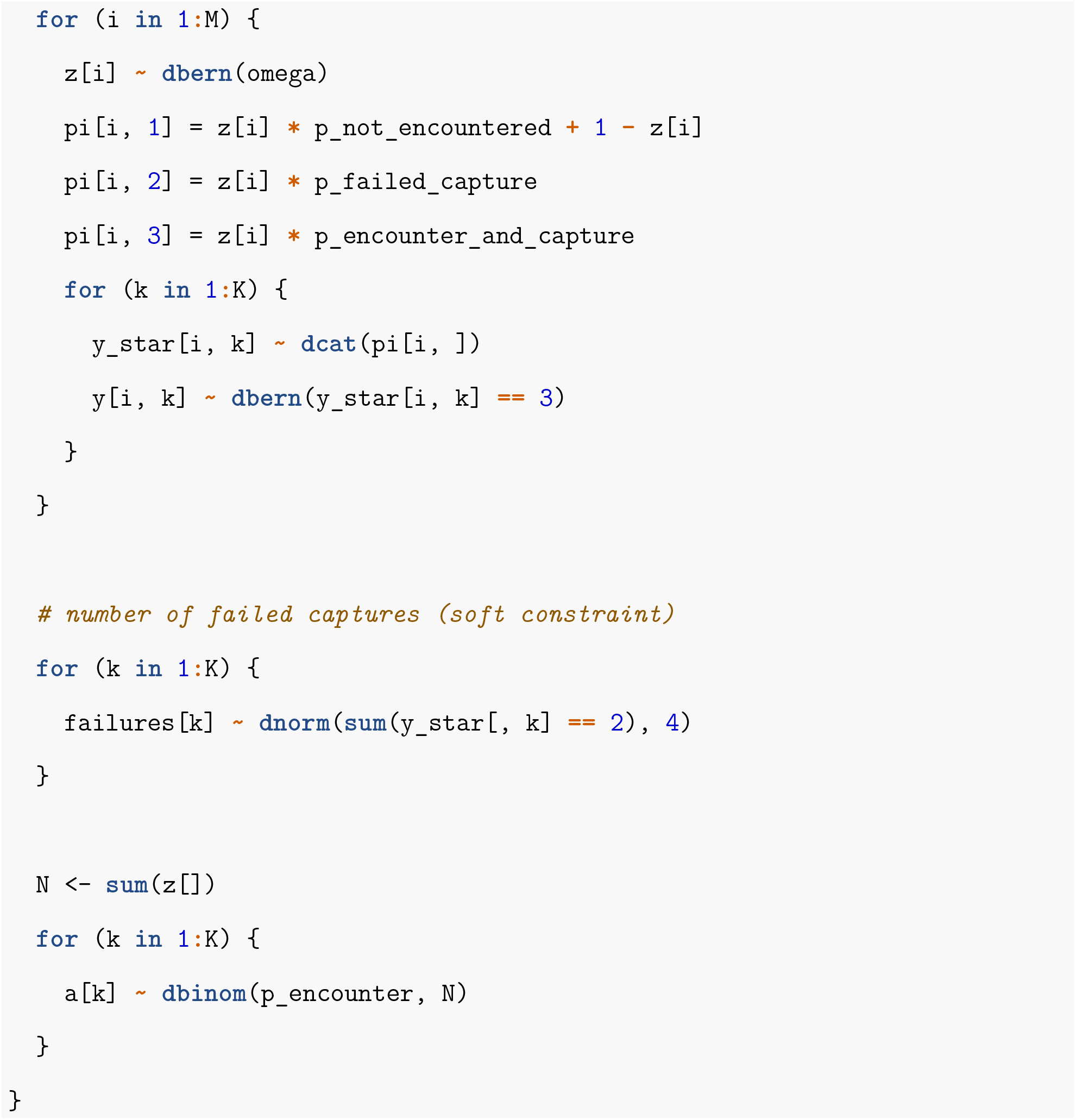

